# A slow unpicking of the landscape thread? Changes in tropical savanna birds on a gradient of habitat modification over 12 years of monitoring

**DOI:** 10.1101/2025.07.29.667582

**Authors:** Alex S. Kutt

## Abstract

**Context:** In south-eastern Australia, land clearing has been extensive, resulting in an avifauna that has been significantly depleted. In northern Australian tropical savannas, land clearing is much less but increasing.

**Aims:** To examine the relative change in bird community composition of a Queensland tropical savanna over a 12-year period where sites represent a gradient of habitat modification. I investigate if these changes trigger avifauna disruption in the wider intact woodlands.

**Methods:** The data were collected from 60 sites sampled seven times between 2004 and 2016, using an eight-count repeat census in each site, in each year of survey. The data was examined via a range of non-parametric multivariate method.

**Key Results:** The analysis of bird community composition indicated that variation was not parallel across the habitat modification gradient (i.e., cleared, thinned sites displayed higher temporal change and instability). The strongest directional change and temporal turnover in β-diversity was strongly linked to the gradient of modification from cleared to thinned, to intact.

**Conclusions:** In this study I demonstrated that, where tree clearing has commenced or trees are thinned (even though it is only a small percentage of the total land cover), this initiates bird community instability, though the surrounding intact vegetation seemed to remain unaffected up until 2016.

**Implications:** Regional changes in bird populations can take many decades to appear after substantial disturbance, and continued monitoring of landscapes being progressively cleared, should be a priority to identify and prevent irreversible collapse of the avifauna, such has occurred in south-eastern Australia.

## Introduction

Despite being the single most significant threat to biodiversity globally (Tilman *et al*. 2017), land clearing for agriculture continues at a depressingly high rate in Australia, and especially in the state of Queensland (Reside *et al*. 2017). This clearing is predominantly broad scale and often including pasture improvement using exotic perennial grasses, targeted clearing of productive areas such as alluvial floodplains for crops, or selective the removal of a proportion of the canopy or understory (i.e., thinning) to promote grass growth (Walker and Weston 1990). The extent of clearing at both a local scale and regional can have small to significant impacts on the fauna (Tilman *et al*. 2017) and in the case of birds - a functionally diverse component of woodlands and forests - the effects can be substantial (Hannah *et al*. 2007; Martin and McIntyre 2007). In southern and eastern Australia, land clearing has been extensive, and resulting in an avifauna that has been significantly depleted and therefore much less resilient to extended periods of climate extremes (Bennett *et al*. 2014). As a consequence, the subtropical and temperate woodland bird community in these areas, has been proposed for listing as a threatened ecological community under national legislation (Fraser *et al*. 2019).

The time lag to species decline and extinction after substantial landscape change, is long and can be commence many decades prior to the depletion of species becoming manifest (Liao *et al*. 2022). In Australia, the ongoing loss and fragmentation of woodlands and forests is linked to the regional loss of species (Ford *et al*. 2009) and the reconfiguration of both the bird community and vegetation has promoted a change in such as the dominance and despotism of hyper-aggressive colonising avifauna that continue to promote the reduction of functional groups such as small-bodied birds (Maron *et al*. 2013). In areas where the clearing for pastoralism is still limited, there is evidence that small scale disturbance can initiate changes in the avifauna (Kutt and Fisher 2011), that could signal the commencement of a longer term impact on birds (Hannah *et al*. 2007).

In this study, I examined the change in bird composition in a tropical savanna woodland in sites sampled over multiple years, and representing a gradient of habitat modification from clearing, thinning and intact vegetation. What is notable in this study is that the landscape is predominantly uncleared, unlike other parts of eastern Australia; however, clearing in Queensland continues to cumulatively increase. I investigated a conceptually straightforward question: what is the change in the bird community composition over the 12-year period of survey, after relative recent (1990-2000s) small scale habitat modification events, and does this trigger on-going community change in the wider landscape (i.e., in both the cleared and thinned vegetation and the adjacent intact vegetation). This paper is a companion piece to a study that modelled in more detail the relationship between individual species abundances and remotely sensed vegetation metrics (Kutt *et al*. 2025c). Articulating how species and communities change in the initial phase of habitat modification is important for planning, managing, and mitigating the long-term negative effects of these changes, especially in the context of overarching climate change.

## Methods

### Study Area

The study area is situated in the Desert Uplands bioregion in central-north Queensland (Fig. 1). The vegetation of the region consists of predominantly *Eucalyptus, Corymbia* and *Acacia* woodlands, open grasslands and ephemeral lakes (DERM 2012). The region is within the southern more arid area of Australia’s tropical savannas, with a long term (1912-2023) mean annual rainfall of 473 mm (Uanda Station, Bureau of Meteorology 2020). The dominant land use is cattle grazing, and vegetation clearing in the Desert Uplands bioregion is estimated to be 7.4%, largely within the Jericho, Bogie River Hills and Prairie – Torrens Creek sub-regions (the latter sub-region being the location of this study) (Queensland Department of Environment and Science 2022).

**Figure 1.**
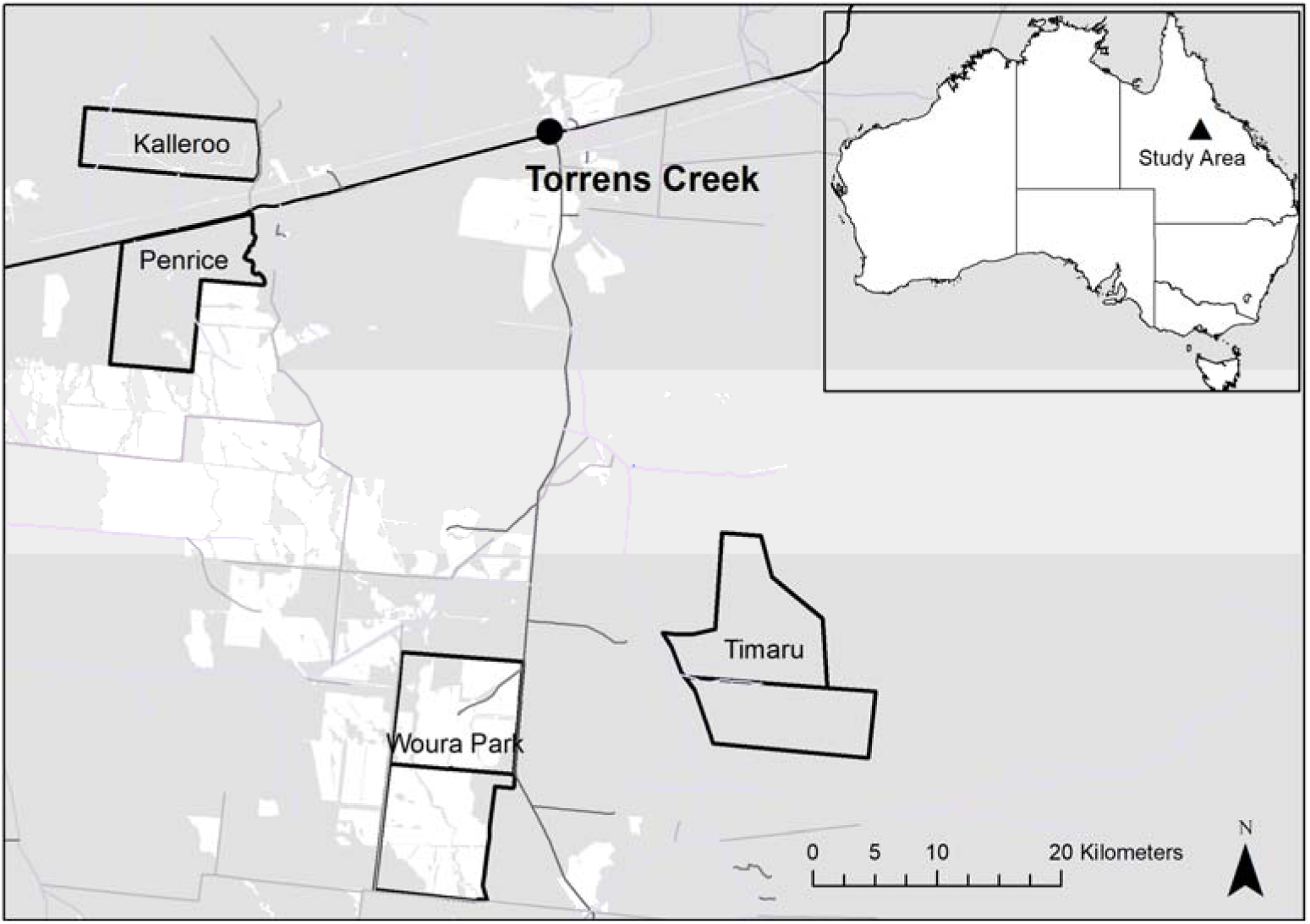
The location of the properties where the bird surveys were undertaken in this study. The grey shaded are indicates intact vegetation and the white areas, cleared. Areas that are thinned are included within the grey shaded areas, as they still represent intact vegetation in the mapping product used for this figure.

The study was undertaken across four properties (Kalleroo, Penrice, Timaru and Woura Park stations) located within a 50 km radius of Torrens Creek (20°48’S, 145°00’E) (Fig. 1). Sixty sites were selected for sampling reflecting three habitat modification types – predominantly cleared of native vegetation, thinned native vegetation, and intact native vegetation (hereafter cleared, thinned and intact). All sites sampled in this study were in a single broad vegetation unit (1: 1,000,000) *Eucalyptus populnea* (poplar box) and *E. melanophloia* (silver-leaved ironbark) or *E. whitei* (White’s ironbark) dry woodlands to open woodlands on sandplains or depositional plains.The three habitat modification types including the thinning method is described in Tassicker *et al*. (2006). The details of the survey periods, and treatment breakdown across the properties is provided (KUTT *et al*. 2025a; Kutt *et al*. 2025c).

### Bird surveys

The bird surveys were undertaken seven times between 2004 and 2016, with no data collected in 2007, 2009-2012 and 2015. All surveys were conducted in the dry season between May and October, the exact timing dictated by access due to prevailing weather and permission from property managers. Sixty sites were sampled in 2004 and 2005 before Kalleroo managers revoked permission to access the property; therefore 50 sites were sampled in the remaining years. One site at Kalleroo could not be accessed in 2005.

The bird surveys were undertaken within a 1 ha site. In each site, 8 x 10-minute meandering diurnal bird census were conducted over a four-day period. Two counts were undertaken at each site on each of the four days: one in the morning between dawn and the three hours after dawn and the other after this period and before dusk. All birds seen and heard within the site were recorded, though species flying over the site were not recorded unless they were interacting with the site (i.e., hunting).

### Analysis

For each site and for each survey, I was able to derive total species richness and total abundance of all birds combined, and abundance of each species. Abundance data were calculated as the total number of birds of each species recorded during each site surveys (i.e., total of eight counts), rather than any calculation of absolute density. Though unadjusted counts can have some biases, the census method used is well-established for Australian tropical savanna bird communities (Woinarski and Tidemann 1991) and has proven to be a reliable index for assessing bird community change in many locations in northern Australia (Hannah *et al*. 2007; Kutt *et al*. 2012b; Kutt *et al*. 2016; Woinarski *et al*. 2012), as well as being a standard state-based approach to bird monitoring (Eyre *et al*. 2018). The scientific and common names, and taxonomic order for birds tabulated, follow the BirdLife Australia working list of Australian birds version 4.1 (BirdLife 2022).

I examined the changes in bird composition across the habitat modification categories and years via several multivariate methods. In the first instance I constructed a site by species abundance array, fourth root transformed the data (to reduce the influence of taxa with high abundance) and then created a Bray-Curtis resemblance (similarity) matrix. I then examined the variation in bird composition across the survey years and habitat modification treatments (and their interaction) via multivariate analysis using the PERMANOVA in PRIMER v7.0.13 (Anderson *et al*. 2008). PERMANOVA is a distance-based, non-parametric, multivariate analysis of variance that calculates a pseudo *F*-statistic and associated *P*-value by means of permutations, rather than relying on normal-theory tables (Anderson *et al*. 2008). In essence this test determines if the centroids (the average point) of different groups (year, habitat modification and the interaction) are significantly different.

Secondly, I examined the homogeneity of multivariate dispersions via PERMDISP (also in PRIMER v7.0.13), which statistically assesses if the spread of data points (sites) is significantly different among groups. I undertook three analyses: for year and habitat modification alone (i.e., the stability of the entire bird community in each year and each habitat category), and then for year and habitat modification categories combined (i.e., 2005 cleared, 2005 thinned, 2005 intact, etc). For visualization, mean distances to centroids were plotted as boxplots for each treatment across all years (intact, thinned, cleared), with jittered points showing site-level variability. The distances to group centroids were calculated using the *betadisper* function in the vegan package (Oksanen *et al*. 2022) for each treatment in each year, providing a measure of within-group multivariate spread. Greater dispersion values indicate higher temporal or spatial heterogeneity in species composition within treatments.

I also compiled the trajectory metrics in two dimensional NMDS to quantify the magnitude and consistency of temporal change via path length (total change through time), net displacement (first–last year distance), and tortuosity (i.e., path length / net displacement). A greater path length and tortuosity indicate larger and more erratic compositional changes (Collins *et al*. 2000). To help visualise these patterns, I created an ordination (non-metric MDS) of the habitat modification and year centroids and overlaid a minimum spanning tree.

I examined temporal β-diversity turnover in bird assemblage composition within each habitat-modification category in each year. For each habitat-modification category, the Bray–Curtis dissimilarity centroid was calculated between the centroid of the baseline year (2004) and the centroid for each subsequent year, providing a measure of compositional divergence through time. The resulting temporal trajectories were plotted for each treatment to visualise changes in assemblage similarity to the baseline. Linear models (*dissimilarity ∼ year*) were fitted separately for each treatment to estimate the rate of compositional change over time (slope of dissimilarity per year), with steeper positive slopes indicating faster divergence and greater temporal turnover. These analyses were undertaken using *vegan* (Oksanen *et al*. 2022) and R 4.4.3 (R Core Team 2024).

To visualise and test how bird assemblage composition in modified habitats changed through time relative to reference conditions, I also used a Principal Response Curve analysis (ter Braak and Šmilauer 2015) using the *prc* function in vegan (Oksanen *et al*. 2022) and R 4.4.3 (R Core Team 2024). PRC is a constrained ordination technique derived from redundancy analysis that extracts the main temporal pattern of treatment effects relative to a specified reference level. In this case, the model structure was specified as *Bird Composition ∼ Habitat Modification × Year*, using the intact sites as the reference treatment, against which trajectories for the thinned and cleared treatments were expressed through time. The PRC curves indicate the deviation of each modification category from the reference across the years, and the proportion of total variance explained by the first PRC axis represents the dominant trajectory of compositional change relative to the reference. PRC also supplies a species weighting vector that indicates how strongly and in which direction individual species contribute to those compositional deviations. High positive values denote species that increased in relative abundance as habitats became more modified (i.e., cleared sites), while negative values identify species that remained largely restricted to intact sites.

Finally for completeness, I examined the variation in species abundance across years in each habitat modification class, using the Kruskal-Wallis non-parametric analysis of variance in Statistica version 13.5.0.17 (TIBCO Software Inc 2018). I tabulated the average abundance (and standard error) for all species for each year and habitat modification category separately, and then indicated which species demonstrated significant variation across each category.

All raw data is available via the scientific data repository *figshare* (Kutt *et al*. 2025b).

## Results

This study recorded a total of 21,899 birds representing 116 species from the 389 sites over the seven years of sampling. The most frequently recorded species were Weebill *Smicrornis brevirostris* (n=341 sites), Singing Honeyeater *Gavicalis virescens* (n=296), Rufous Whistler *Pachycephala rufiventris* (n=282), Striated Pardalote *Pardalotus striatus* (n=282) and Yellow-throated Miner *Manorina flavigula* (n=265). The most abundantly recorded species were Weebill (n=3263 records), Budgerigar *Melopsittacus undulatus* (n=1400), Variegated Fairy-wren *Malurus lamberti* (n=1161), Zebra Finch *Taeniopygia castanotis* (n=1090) and Yellow-throated Miner (n= 872). Twelve bird species were only recorded once, and 10 of these were only via a single record. Considering the distribution of species across the habitat modification categories, 23 species were only recorded in a single category, while 70 species were recorded in all three; for years, this breakdown was 15 in a single year and 48 in every year.

For all sites combined, the highest mean site species abundance was in 2005, and highest mean site species richness was in 2014 (Table 1). Considering cleared sites alone, highest abundance was in 2005, and highest species richness in 2013, for thinned sites alone, highest abundance was in 2005, and highest species richness in 2014 and for intact sites, highest abundance was also in 2005, and highest species richness in 2014 (Table 1).

**Table 1.**
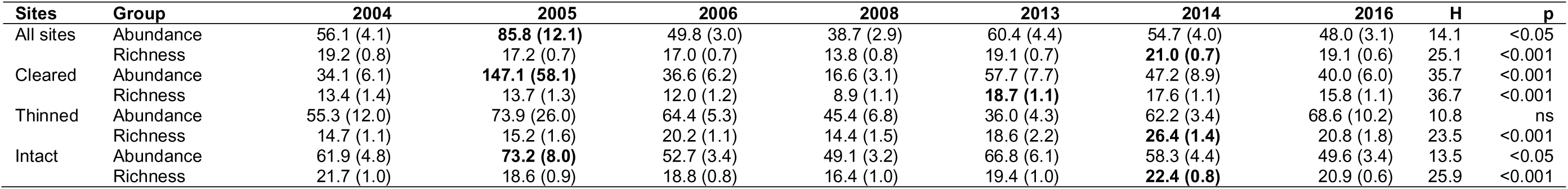
The mean (and standard error) of bird abundance and species richness recorded across all surveys for each of the years. Data presented is all sites combined and then for each habitat modification category.

Variation in individual species abundance for each habitat modification category across the years is presented in Supplementary Table 1. As this is not the direct focus of the manuscript, these results are not considered further, except to note that in cleared sites 41 species indicated significant variation in abundance across the years of sampling (from a total of 101 species recorded), in thinned sites 22 species indicated significant variation in abundance across the years of sampling (from a total of 85 species recorded), and in intact sites 55 species indicated significant variation in abundance across the years of sampling (from a total of 110 species recorded).

The PERMANOVA results indicated that there was a strong overall effect of year (df = 6, Pseudo-F = 6.94, P(perm) = 0.001), habitat modification (df = 2, Pseudo-F = 25.4, P(perm) = 0.001) and the interaction (df = 20, Pseudo-F = 2.43, P(perm) = 0.001) on bird community composition. The larger Pseudo-F (a measure of effect size) for habitat modification suggests this is a stronger effect than year and the interaction. The PERMDISP indicated that was a strong deviation from the centroid (i.e., evidence of dispersion) for habitat modification, (F = 47.3, df1 = 2, df2 = 386, P(perm): 0.001) but less so for year (F = 10.0, df1 = 6, df2 = 382 P(perm) = 0.001) and the interaction (F = 11.3, df1 = 20, df2 = 368 P(perm) = 0.001). The boxplots of the multivariate dispersion provide further detail of the pattern (Fig. 2), namely, that intact sites consistently showed the lowest distances to group centroids, indicating relatively homogeneous and stable assemblage composition within years, whereas thinned sites exhibited intermediate levels of dispersion, with modest year-to-year fluctuations and cleared sites displayed the greatest within-group dispersion across most years, reflecting higher spatial or temporal heterogeneity in community composition.

**Figure 2.**
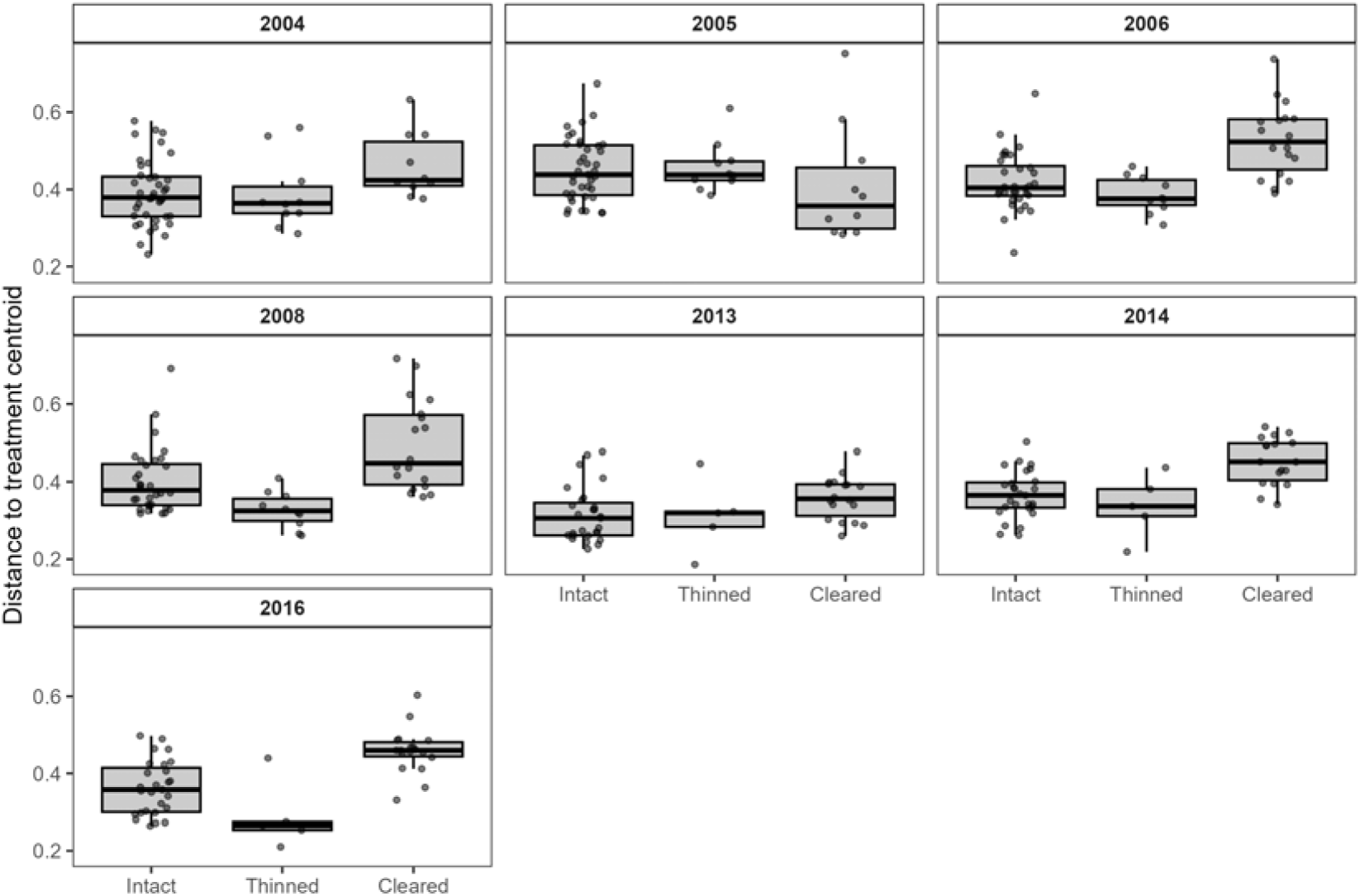
Boxplots of multivariate dispersion (Bray–Curtis distance to the group centroid) for bird assemblages under three habitat-modification treatments (intact, thinned, and cleared) across survey years. Points denote individual site distances to the treatment centroid.

In NMDS space, cleared sites exhibited the greatest total path length (1.64), followed by thinned (1.35) and intact (0.70), indicating the largest cumulative change in cleared habitats. However, tortuosity, reflecting irregular or non-directional change, was highest in Thinned (38.4), compared with Cleared (8.52) and Intact (6.11). This pattern suggests that thinned assemblages changed most erratically, whereas intact communities were the most stable through time (Fig. 3).

**Figure 3.**
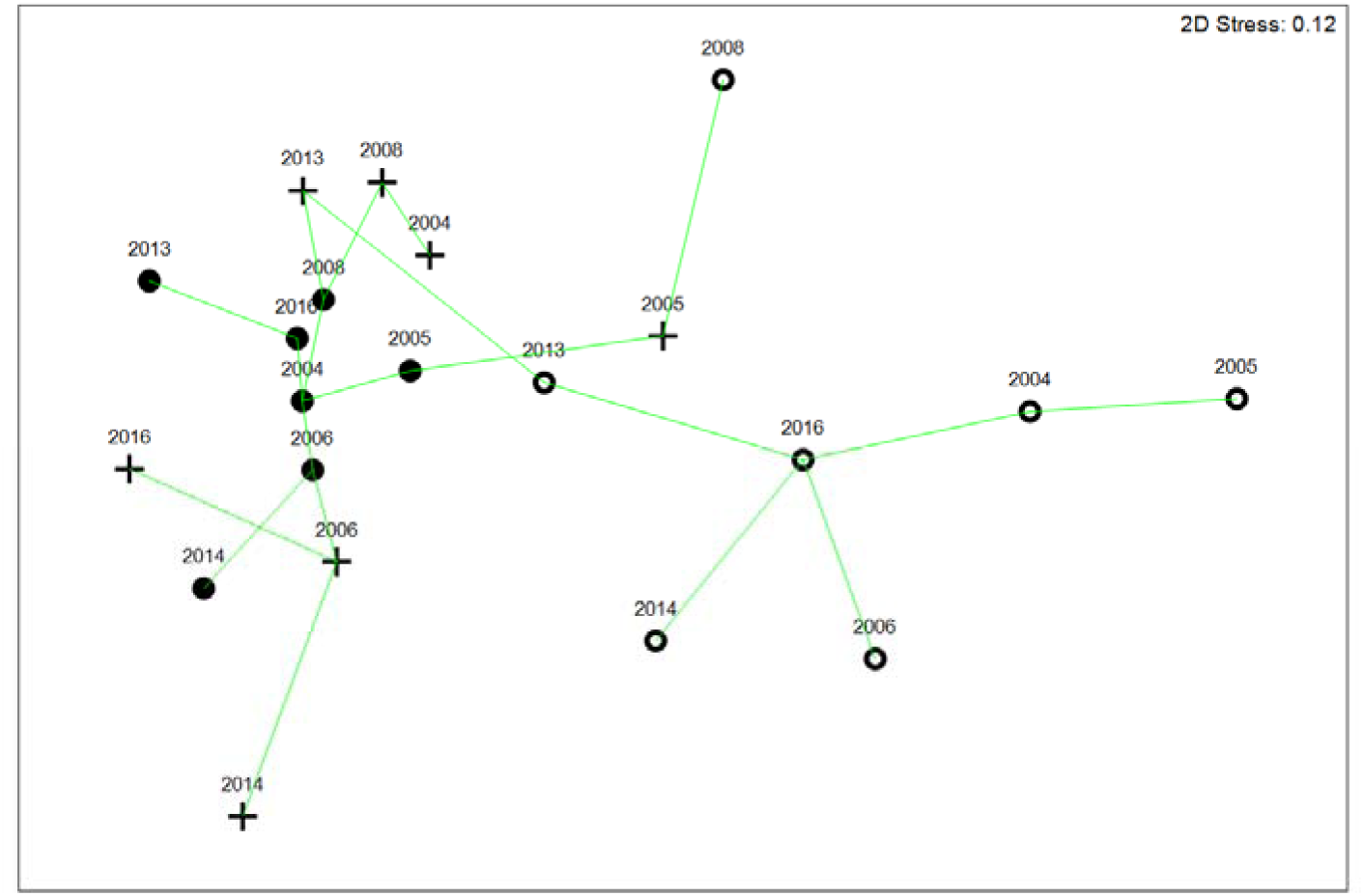
Ordination via non-metric multidimensional scaling indicating the centroids for bird composition for each habitat modification category by year, joined by a minimum spanning tree. Cleared sites are the open circles (o), thinned sites are the crosses (+) and intact sites are solid circles (●).

Temporal β-diversity analyses revealed clear differences in the rate of compositional change among habitat-modification treatments (Fig. 4). Bird assemblages in cleared sites exhibited the greatest and most consistent increase in Bray–Curtis dissimilarity from baseline conditions through time, indicating progressive and substantial community turnover. The thinned sites showed intermediate levels of change, with moderate increases in dissimilarity suggesting ongoing but less pronounced compositional shifts. Finally, the intact habitats remained relatively stable across survey years, with only minor departures from baseline assemblages. Linear regression slopes of temporal β-diversity against year confirmed this pattern. The strongest positive slope occurred in cleared sites (Δ = 0.0487 yr⁻¹, *p* ≈ 0.06, R² = 0.54), followed by thinned (Δ = 0.0322 yr⁻¹, *p* ≈ 0.07, R² = 0.51) and intact (Δ = 0.0370 yr⁻¹, *p* ≈ 0.27, R² = 0.24).

**Figure 4.**
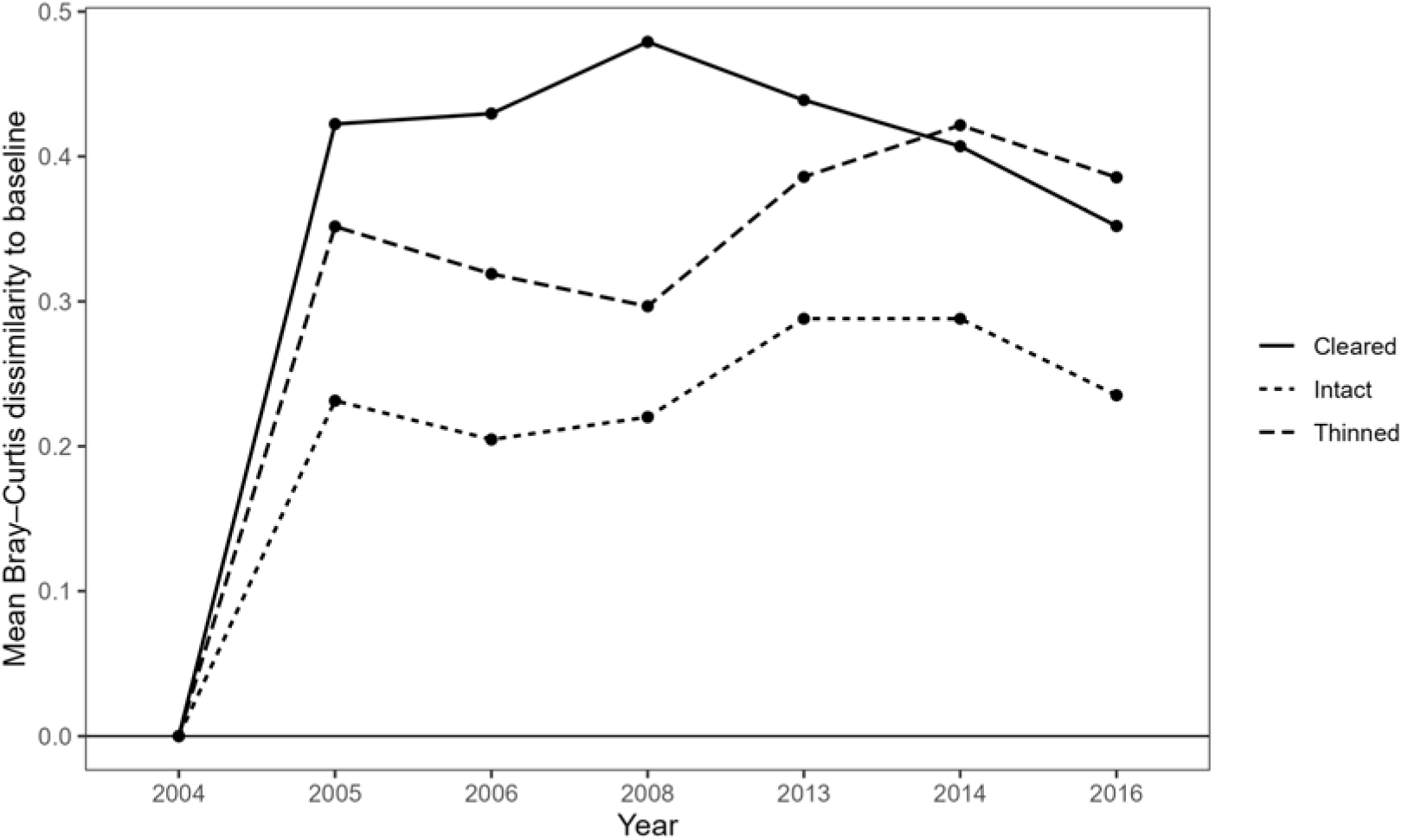
Temporal trajectories of bird assemblage change, expressed as mean Bray–Curtis dissimilarity from the baseline year for each habitat-modification treatment (intact, thinned, cleared). Lines represent treatment-specific community turnover through time, with higher values indicating greater compositional divergence from baseline assemblages. Points show annual mean dissimilarities, and the horizontal line at zero indicates no change relative to baseline.

The overall PRC model was significant (*F* = 4.12, *p* = 0.001; 999 permutations), indicating that bird community composition in modified treatments changed markedly and directionally through time relative to intact habitats. The first PRC axis accounted for most of the habitat modification related temporal variance, indicating a clear separation between the intact reference data and the two modified habitats (Fig. 5). Bird community composition in cleared habitats exhibited the greatest and most consistent deviation from intact bird community baseline, reflecting progressive compositional change over the years. The thinned sites habitats showed intermediate divergence, with less fluctuation and intact habitats remained close to the baseline, indicating high temporal stability in community composition. The species weights (greatest magnitude, either positive or negative) indicated species characteristic of the gradient from the intact to modified (cleared and thinned sites) were typical of good condition woodlands (e.g., Grey-crowned Babbler, Grey Shrike-thrush, Spiny-cheeked Honeyeater, Brown Treecreeper, Striated Pardalote, Rufous Whistler) and disturbed or cleared woodlands (e.g., Zebra Finch, Galah, Black-faced Woodswallow, Australian Magpie, Brown Falcon, Crested Pigeon) (Fig. 6).

**Fig. 5.**
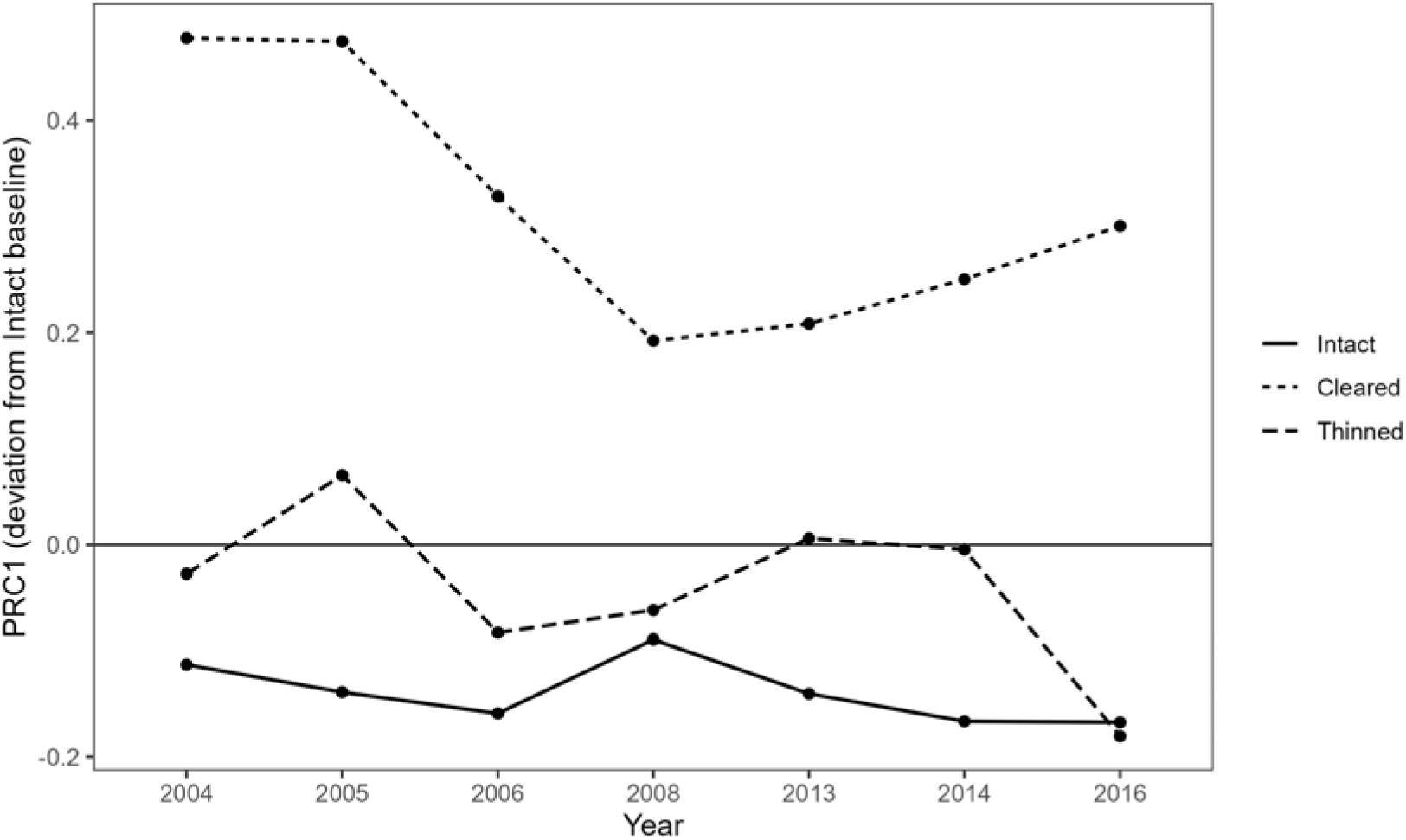
Principal Response Curves (PRC) showing treatment-specific temporal trajectories of bird assemblage composition relative to intact reference habitats. The horizontal zero line represents no difference from the reference condition, with greater deviations indicating stronger compositional divergence.

**Figure 6.**
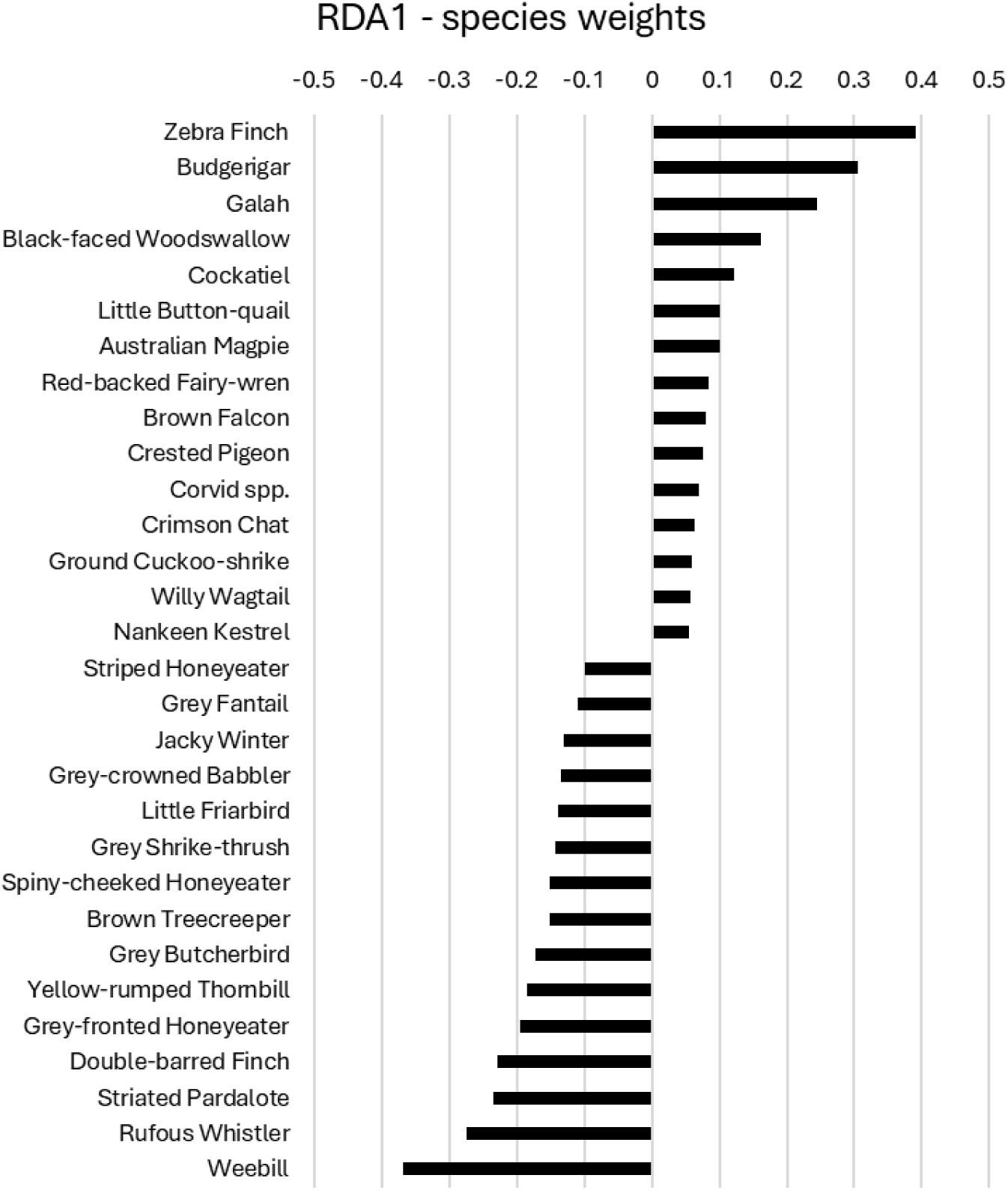
Species weights from the Principal Response Curve analysis showing the taxa most strongly contributing to compositional differences among habitat-modification treatments relative to the intact baseline. The top 15 species characteristic of intact sites and the top 15 species characteristic of the most modified sites (cleared) are presented. Larger absolute weights indicate taxa exerting greater influence on treatment separation and temporal community trajectories. Positive values denote species that increased in relative abundance as habitats became more modified, while negative values identify species that declined or remained largely restricted to intact conditions.

## Discussion

The clearing of native vegetation in many parts of the world, including north-eastern and south-eastern Australia, have caused a substantial past and continuing disarray of bird communities and species (Berryman *et al*. 2024; Thomson *et al*. 2015). The primary cause of bird community decline in Australia is land clearing associated with agriculture (Lindenmayer 2022b). In this study I examined the pattern of bird species in a series of sites (sampled over multiple years), representing the preliminary stages of more extensive land clearing that has already occurred elsewhere in Australia. In general, there were well-defined differences in species composition across the cleared and intact tropical savanna woodlands, and more muted intermediate differences with the thinned vegetation. The concept of land clearing thresholds for the prevention of species decline and local extirpation have been well articulated (McAlpine *et al*. 2002), yet the attrition of native vegetation continues in a dispiritingly substantial manner on many land tenures with little effective legislative oversight (Ward *et al*. 2019). In this study I found that small scale clearing and thinning triggers a degree of instability in the bird community in the cleared sites, which may be the precursor of long-term more permanent depletion of the birds in the modified habitat, even when the surrounding matrix is still intact. The data did not indicate a change in the adjacent intact vegetation, but it is conceivable that over time (i.e., decades) this disruption will influence the surrounding vegetation, for example via edge effects, exotic pasture invasion or changes in stocking rates.

I identify a few limitations to the study and analysis. Firstly, as each site in each year, was censused eight times, occupancy analysis may have provided more detailed and revealing data regarding the changes in bird species over time in each modification category (Montague-Drake *et al*. 2009). However, over the years of survey, the data was not consistently structured in the data base (i.e., summaries versus each count separately) to allow this analysis. Secondly, even though the study spanned 12 years, the sites were not annually monitored, which provides some gaps the annual patterns, and any evidence of unusual or predictable changes from year to year. Lastly, site based environmental factors, including changes to the site management of these pastoral properties (i.e., stocking rates, exotic pasture incursion, regrowth or re-clearing), which does not allow for a more nuanced understanding of the mechanisms of change in the bird communities across the sites and habitat modification categories. Any future work that may revisit these sites, should attempt to address these issues. Regardless, this study provides a unique insight into the changes over a 12-year period of a bird community at the initial phase of disturbance.

The analysis of bird community composition change indicated that the variation was not parallel across the three habitat modification treatments (i.e., the significant interacting effect in the PERMANOVA). Clearing and thinning have a strong effect on the avifauna in the study area, but the temporal changes differed on the gradient of modification. Though heterogenous landscapes can support multiple bird communities and increase the bird diversity across the wider matrix, woodland and forest areas are numerically the most significant for bird conservation (Lu *et al*. 2024). However the species in cleared areas are not necessarily equivalent to natural grasslands, despite species overlap, and this may be in part of changes in resources, such as available diet (Houston *et al*. 2023) and differences in ground cover. In general, when land clearing continues, the bird depletion continues (Lindenmayer 2022a) and only woodland restoration can reverse this effect (Bennett *et al*. 2022). In this study I found that disturbance increases community heterogeneity, but with less stability in the modified sites. Evidence from the sites further south to the study area with a greater extent of landscape modification and fragmentation (Hannah *et al*. 2007) indicated that bird species richness in cleared sites were substantially reduced (8.1 species per site compared to this study 14.4 [average over all years combined]) whereas species richness in intact sites was remarkably coincident (19.9 species versus 19.8 in this study[average over all years combined]).

The bird assemblages underwent the strongest directional change through time in the cleared sites, with a slight trajectory towards the intact sites in 2008, which then reversed. Thinned sites remained closer to the intact sites. This pattern is driven by disturbance-tolerant taxa, or species that fluctuate with rainfall events (the study sites being on the edge of more arid bioregion). Woodland bird assemblages in Australia can be highly variable from year to year (Maron *et al*. 2005), and these patterns can be driven by climatic events (Bennett *et al*. 2014), associated resource pulses or declines (Collett *et al*. 2022) as well as seasonal regional or continental migration patterns (Chan 2001). The species that were characteristic of, and influential in the temporal trajectories of cleared were granivores that can demonstrate distinct pulses of abundance (i.e., Little Button-quail, Galah, Budgerigar, Zebra Finch), species that typically occur in cleared and disturbed habitat (i.e., Crested Pigeon, Australian Magpie), and species that occur in natural grasslands (i.e., Australian Kestrel, Ground Cuckoo-shrike, Black-faced Woodswallow). Conversely the species in the intact sites were characteristic of intact and good condition woodlands (Fraser *et al*. 2019). This contrast is typical of those occurring across a treed to treeless gradient, such as woodland dependent birds (i.e., species in Meliphagidae, Pachycephalidae, Acanthizidae) and species associated with grasslands and bare ground (Barrett *et al*. 1994) and the woodland-grassland nexus in tropical savannas (Kutt and Fisher 2011; Kutt *et al*. 2012a; Woinarski *et al*. 2000; Woinarski and Tidemann 1991).

The temporal turnover in β-diversity in the study sites indicated that this pattern was strongly linked to the gradient of modification from cleared to thinned, to intact. In addition, there are large directional community shifts in the cleared sites (and a lesser degree in thinned sites) compared to the intact sites which are relatively stable. Disturbance driven variability can be an important precursor to more persistent ecological change (Fraterrigo and Rusak 2008), and though variability can be natural, driven by, for example, climatic fluctuations (Recher and Davis 2014), vegetation changes can have a stronger effect on bird composition compared to weather patterns (Pavey and Nano 2009). The mechanisms for these species decline in south-eastern Australia, where there is more substantial habitat loss and fragmentation, include reduced dispersal and higher nest predation and then reduced breeding leading to local extirpation (Ford *et al*. 2009). Though in this study I did not have site based environmental data to demonstrate the oscillation in bird community is linked to an equivalent fluctuation in, and possible reduction in, key resources for species, there is ample evidence, albeit on a larger scale of clearing, that once clearing commences, this unlocks a cascading influence on both resources, threats and habitat conditions, that create an eventual species deficit (Ford *et al*. 2009).

Changes in the vertebrate fauna of tropical savannas in northern Australia are evident, which mostly have become manifest in the mammal fauna (Fisher *et al*. 2014). The data for bird assemblages seem to suggest more resilience poor land management (Woinarski *et al*. 2012), though there is likely a geographical effect. In some areas of north-eastern Australia where overgrazing has significantly altered intact vegetation (i.e., permanently altered the ground cover, and removed the shrub layer), the avifauna can also be depleted (Kutt and Fisher 2011). In this study I have demonstrated that, where tree clearing has commenced or trees are thinned (even though it is only a small percentage of the total land cover), this seems to initiate instability in these modified habitats. During the timespan of this study, the intact bird communities did not follow this pattern of high variability. Regional changes in bird populations can take many decades to appear after substantial disturbance (i.e., extinction debt Szabo *et al*. 2011), though at a landscape scale where change is multi-directional or more limited, these can be both debits and credits (Haddou *et al*. 2022). The sites described in this study were last sampled in 2016, and surveys have been discontinued. Long term monitoring is a critical action for improved landscape management and conservation (Lindenmayer *et al*. 2022). In this study I present data that hints at an unpicking of the landscape condition for woodland birds; extension, enhancement or even recasting of long-term research, such as those presented here, should be ongoing, and include more specific focus on biological mechanisms for species variation, and if, over time, the adjacent intact vegetation also succumbs to gradual disruption of the woodland bird community.

## Acknowledgements

I am grateful for the help of landholders in granting us access to their properties at Woura Park, Timaru, Penrice and Kalleroo, and the many people who assisted in the surveys, from the Australian CSIRO and Queensland Department of Environment and Science. This project was supported by the Queensland Government and the Australian Government (CSIRO, National Heritage Trust, Tropical Savanna Cooperative Research Centre, TERN). All surveys were conducted under the Queensland Government Scientific Purposes Permit number WITK04645707.

## Notes

https://doi.org/10.26188/29710859.v1

### Competing Interest Statement

The authors have declared no competing interest.

### Summary of Updates

Following recommendations from referee reports, the data analysis has been updated to focus more on community composition change, in the different habitat modification categories, across the years of survey, rather than a focus on changes over years alone, and modification categories. New multivariate analyses are included, such as PERMANOVA, PermDisp, Principal Response Curves and beta-diversity turnover. Some of the Discussion has changed as a consequence.

