## Supplemental Table 1 for "A slow unpicking of the landscape thread? Changes in tropical savanna birds on a gradient of habitat modification over 12 years of monitoring"

**Supplementary Table 1.** The mean (and standard error) of bird abundance and species richness, and each species, recorded across all surveys, for each of the years. Significant variation across the years is tested via Kruskal-Wallis non-parametric analysis of variance. H is the test statistic and p-values indicate three significance levels, and ns (not significant).

| **Bird species** | **2004** | **2005** | **2006** | **2008** | **2013** | **2014** | **2016** | **H** | **p value** |
| --- | --- | --- | --- | --- | --- | --- | --- | --- | --- |
| **CLEARED** | | | | | | | | | |
| Emu |  |  | 0.111 (0.111) | 0.056 (0.056) |  | 0.056 (0.056) | 0.056 (0.056) | 2.2 | ns |
| Brown Quail |  |  | 0.167 (0.167) |  |  |  |  | 5.1 | ns |
| Common Bronzewing |  |  | 0.333 (0.243) | 0.167 (0.090) | 0.167 (0.090) | 0.278 (0.158) | 0.611 (0.397) | 4.7 | ns |
| Crested Pigeon | 0.700 (0.396) |  | 0.222 (0.173) | 0.167 (0.090) | 0.389 (0.118) | 0.667 (0.198) | 0.222 (0.101) | 14.3 | <0.05 |
| Southern Squatter Pigeon |  |  |  |  |  |  | 0.056 (0.056) | 5.1 | ns |
| Diamond Dove |  |  | 0.167 (0.121) |  |  |  |  | 10.3 | ns |
| Peaceful Dove | 0.100 (0.100) |  | 0.167 (0.121) |  | 0.222 (0.101) | 0.333 (0.114) | 0.056 (0.056) | 13.0 | <0.05 |
| Bar-shouldered Dove |  |  |  |  | 0.111 (0.076) | 0.056 (0.056) |  | 7.3 | ns |
| Australian Owlet-nightjar |  | 0.100 (0.100) |  |  |  |  |  | 10.0 | ns |
| White-necked Heron | 0.100 (0.100) |  |  |  |  |  |  | 10.0 | ns |
| Black-breasted Buzzard | 0.100 (0.100) |  |  | 0.056 (0.056) |  |  |  | 6.6 | ns |
| Whistling Kite |  |  |  | 0.056 (0.056) | 0.056 (0.056) | 0.056 (0.056) |  | 3.2 | ns |
| Black Kite |  | 0.200 (0.133) |  | 0.056 (0.056) |  |  | 0.222 (0.129) | 12.1 | ns |
| Brown Goshawk | 0.200 (0.133) |  |  |  |  |  |  | 20.2 | <0.01 |
| Collared Sparrowhawk |  |  |  |  |  | 0.056 (0.056) |  | 5.1 | ns |
| Spotted Harrier |  |  | 0.111 (0.111) |  |  |  | 0.111 (0.076) | 7.2 | ns |
| Wedge-tailed Eagle | 0.200 (0.133) | 0.100 (0.100) | 0.056 (0.056) | 0.111 (0.076) | 0.056 (0.056) |  | 0.278 (0.135) | 7.0 | ns |
| Little Eagle |  |  |  |  |  | 0.056 (0.056) |  | 5.1 | ns |
| Nankeen Kestrel | 0.300 (0.153) | 0.200 (0.200) | 0.056 (0.056) | 0.056 (0.056) | 0.111 (0.076) | 0.056 (0.056) | 0.222 (0.129) | 5.8 | ns |
| Brown Falcon | 0.900 (0.233) | 0.700 (0.213) | 0.111 (0.076) | 0.278 (0.109) |  | 0.278 (0.135) | 0.611 (0.143) | 28.6 | <0.001 |
| Australian Hobby |  | 0.200 (0.133) | 0.056 (0.056) |  | 0.056 (0.056) |  |  | 10.3 | ns |
| Australian Bustard | 0.100 (0.100) | 0.200 (0.133) |  | 0.056 (0.056) | 0.111 (0.076) | 0.056 (0.056) | 0.444 (0.202) | 9.7 | ns |
| Little Button-quail | 0.600 (0.400) | 3.900 (1.059) | 0.444 (0.217) |  |  |  | 0.611 (0.413) | 51.8 | <0.001 |
| Red-tailed Black-Cockatoo |  |  |  |  | 0.056 (0.056) | 0.111 (0.076) |  | 7.3 | ns |
| Galah | 0.800 (0.416) | 22.500 (22.059) | 2.278 (1.255) |  | 4.500 (2.231) | 1.500 (0.390) | 1.167 (0.336) | 23.1 | <0.001 |
| Sulphur-crested Cockatoo |  |  |  |  | 0.056 (0.056) | 0.111 (0.076) | 0.056 (0.056) | 5.3 | ns |
| Cockatiel | 0.100 (0.100) | 2.700 (1.334) | 1.167 (0.500) |  | 1.000 (0.662) | 3.778 (2.914) | 0.889 (0.604) | 19.1 | <0.01 |
| Rainbow Lorikeet |  |  |  | 0.056 (0.056) | 0.556 (0.305) | 0.056 (0.056) | 0.167 (0.090) | 10.3 | ns |
| Red-winged Parrot | 0.100 (0.100) |  | 0.722 (0.609) |  | 0.167 (0.121) | 0.944 (0.488) | 0.333 (0.140) | 11.9 | ns |
| Pale-headed Rosella |  |  | 0.611 (0.421) |  | 0.722 (0.195) | 0.444 (0.185) | 0.222 (0.129) | 22.8 | <0.001 |
| Budgerigar | 2.300 (1.415) | 41.100 (27.048) | 2.667 (1.515) | 0.056 (0.056) | 9.611 (4.103) | 7.944 (5.728) | 1.167 (0.601) | 37.5 | <0.001 |
| Horsfield's Bronze-Cuckoo |  |  | 0.444 (0.145) |  | 0.111 (0.076) | 0.056 (0.056) | 0.222 (0.222) | 21.2 | <0.01 |
| Shining Bronze-Cuckoo |  |  |  |  | 0.056 (0.056) |  |  | 5.1 | ns |
| Pallid Cuckoo |  |  | 0.056 (0.056) |  |  |  | 0.056 (0.056) | 4.1 | ns |
| Southern Boobook |  | 0.100 (0.100) |  |  |  |  |  | 10.0 | ns |
| Blue-winged Kookaburra |  |  |  |  |  |  | 0.111 (0.076) | 10.3 | ns |
| Red-backed Kingfisher | 0.100 (0.100) | 0.300 (0.213) | 0.500 (0.259) | 0.111 (0.076) |  |  | 0.056 (0.056) | 9.5 | ns |
| Rainbow Bee-eater |  |  | 0.278 (0.135) |  |  | 0.833 (0.493) |  | 27.5 | <0.001 |
| Brown Treecreeper |  |  | 0.056 (0.056) | 0.056 (0.056) |  | 0.056 (0.056) |  | 3.2 | ns |
| Spotted Bowerbird |  |  |  |  | 0.056 (0.056) |  | 0.111 (0.111) | 4.1 | ns |
| Great Bowerbird |  |  |  | 0.056 (0.056) |  |  |  | 5.1 | ns |
| unknown Fairy-wren |  |  |  |  |  | 0.056 (0.056) | 0.389 (0.231) | 11.7 | ns |
| Red-backed Fairy-wren |  |  | 0.333 (0.243) |  | 2.056 (0.931) | 2.111 (0.863) | 2.500 (1.253) | 21.9 | <0.01 |
| White-winged Fairy-wren |  |  |  |  |  |  | 0.278 (0.177) | 15.6 | <0.05 |
| Variegated Fairy-wren | 2.300 (1.065) | 1.000 (0.537) | 1.000 (0.333) | 0.500 (0.336) | 2.389 (0.954) | 3.000 (0.828) | 3.444 (0.994) | 15.1 | <0.05 |
| Weebill | 1.200 (0.786) | 3.400 (1.157) | 2.000 (0.686) | 4.222 (1.201) | 11.222 (1.507) | 2.056 (0.431) | 2.278 (1.182) | 33.0 | <0.001 |
| Western Gerygone |  |  | 0.056 (0.056) |  | 0.167 (0.090) | 0.056 (0.056) |  | 8.8 | ns |
| White-throated Gerygone |  |  | 0.056 (0.056) |  |  |  |  | 5.1 | ns |
| Yellow-rumped Thornbill | 0.300 (0.300) | 0.200 (0.200) | 1.667 (1.057) | 0.944 (0.545) | 0.944 (0.400) | 0.333 (0.140) | 0.667 (0.333) | 5.5 | ns |
| Buff-rumped Thornbill | 0.100 (0.100) |  |  |  |  |  | 0.056 (0.056) | 6.6 | ns |
| Inland Thornbill |  |  | 0.444 (0.271) |  | 0.278 (0.226) | 0.056 (0.056) | 0.389 (0.293) | 6.2 | ns |
| Red-browed Pardalote |  |  |  |  | 0.278 (0.109) | 0.056 (0.056) |  | 21.5 | <0.01 |
| Striated Pardalote | 0.800 (0.416) | 0.800 (0.291) | 0.667 (0.412) | 1.222 (0.461) | 1.889 (0.464) | 0.222 (0.129) | 0.222 (0.101) | 26.6 | <0.001 |
| Singing Honeyeater | 1.900 (0.586) | 0.900 (0.314) | 1.389 (0.380) | 0.500 (0.146) | 2.889 (0.511) | 2.222 (0.417) | 1.167 (0.232) | 26.0 | <0.001 |
| Grey-fronted Honeyeater |  |  | 0.167 (0.121) | 0.167 (0.167) | 0.056 (0.056) |  | 0.056 (0.056) | 3.7 | ns |
| Yellow-throated Miner | 1.300 (0.597) | 1.300 (0.559) | 0.889 (0.457) | 0.611 (0.183) | 1.833 (0.305) | 1.333 (0.443) | 1.056 (0.318) | 13.2 | <0.05 |
| Spiny-cheeked Honeyeater | 0.200 (0.133) | 0.100 (0.100) | 0.167 (0.167) |  | 0.333 (0.114) | 0.222 (0.129) |  | 13.9 | <0.05 |
| Rufous-throated Honeyeater |  |  |  |  |  | 0.056 (0.056) |  | 5.1 | ns |
| Crimson Chat |  | 2.600 (0.833) |  | 0.611 (0.421) | 2.056 (1.259) |  |  | 41.7 | <0.001 |
| Blue-faced Honeyeater |  |  |  |  |  |  | 0.111 (0.076) | 10.3 | ns |
| Noisy Friarbird |  |  | 0.111 (0.076) |  |  | 0.111 (0.076) |  | 8.5 | ns |
| Little Friarbird | 0.100 (0.100) | 0.100 (0.100) | 0.278 (0.158) | 0.056 (0.056) |  | 1.278 (0.441) |  | 28.0 | <0.001 |
| Striped Honeyeater |  |  |  | 0.056 (0.056) | 0.167 (0.090) | 0.111 (0.076) | 0.111 (0.111) | 6.3 | ns |
| Grey-crowned Babbler |  |  | 0.556 (0.364) | 0.222 (0.173) | 1.056 (0.286) | 0.667 (0.388) | 1.667 (0.828) | 21.1 | <0.01 |
| Varied Sittella |  | 0.800 (0.800) | 0.056 (0.056) | 0.278 (0.158) | 0.333 (0.229) | 0.444 (0.246) | 0.778 (0.440) | 4.1 | ns |
| Ground Cuckoo-shrike | 0.600 (0.499) | 0.200 (0.133) | 0.111 (0.076) |  | 0.278 (0.177) | 0.056 (0.056) | 0.500 (0.232) | 8.0 | ns |
| Black-faced Cuckoo-shrike | 0.600 (0.163) | 0.300 (0.213) | 2.000 (0.667) | 0.056 (0.056) | 0.389 (0.143) | 0.444 (0.145) | 0.611 (0.244) | 19.6 | <0.01 |
| White-bellied Cuckoo-shrike |  |  |  |  | 0.056 (0.056) |  |  | 5.1 | ns |
| White-winged Triller | 0.200 (0.133) | 1.600 (1.222) | 1.056 (0.424) |  | 0.222 (0.173) | 0.333 (0.162) |  | 15.4 | <0.05 |
| Rufous Whistler | 0.500 (0.269) | 0.300 (0.300) | 0.722 (0.411) | 0.722 (0.360) | 1.333 (0.589) | 0.778 (0.222) | 0.556 (0.202) | 10.1 | ns |
| Grey Shrike-thrush | 0.100 (0.100) |  |  | 0.222 (0.101) | 0.222 (0.173) | 0.167 (0.090) | 0.167 (0.121) | 6.1 | ns |
| Crested Bellbird | 0.200 (0.133) |  | 0.778 (0.173) | 0.167 (0.090) | 0.889 (0.111) | 0.778 (0.152) | 0.944 (0.235) | 32.6 | <0.001 |
| Olive-backed Oriole |  |  |  |  | 0.056 (0.056) | 0.167 (0.090) | 0.056 (0.056) | 8.8 | ns |
| White-breasted Woodswallow |  | 0.400 (0.267) |  |  |  |  |  | 20.2 | <0.01 |
| Masked Woodswallow |  | 0.200 (0.200) | 0.222 (0.152) |  |  | 0.333 (0.114) | 0.056 (0.056) | 16.8 | <0.05 |
| White-browed Woodswallow |  |  |  |  |  |  | 0.222 (0.129) | 15.6 | <0.05 |
| Black-faced Woodswallow | 1.600 (0.909) | 4.000 (1.483) | 1.833 (0.648) | 0.889 (0.212) | 0.778 (0.392) | 0.889 (0.342) | 2.000 (0.691) | 10.6 | ns |
| Little Woodswallow | 0.300 (0.153) | 0.200 (0.200) | 0.222 (0.173) | 0.056 (0.056) |  | 0.278 (0.177) |  | 10.1 | ns |
| Grey Butcherbird |  |  | 0.167 (0.090) | 0.389 (0.118) | 0.500 (0.121) | 0.278 (0.135) | 0.167 (0.090) | 15.9 | <0.05 |
| Pied Butcherbird | 0.500 (0.307) | 0.300 (0.153) | 0.389 (0.183) | 0.444 (0.185) | 1.167 (0.146) | 0.944 (0.235) | 1.278 (0.226) | 31.0 | <0.001 |
| Australian Magpie | 0.900 (0.277) | 0.800 (0.512) | 0.333 (0.162) | 0.778 (0.101) | 1.056 (0.151) | 0.667 (0.114) | 1.056 (0.221) | 17.0 | <0.01 |
| Grey Fantail | 0.100 (0.100) |  | 0.056 (0.056) | 0.222 (0.152) | 0.778 (0.329) |  | 0.278 (0.177) | 13.2 | <0.05 |
| Willie Wagtail | 1.500 (0.687) | 0.600 (0.306) | 0.333 (0.162) | 0.611 (0.183) | 1.111 (0.290) | 0.444 (0.121) | 0.722 (0.211) | 8.7 | ns |
| Crow / Raven | 1.900 (1.140) | 2.500 (1.790) |  |  |  | 0.556 (0.121) |  | 51.1 | <0.001 |
| Torresian Crow |  |  | 0.667 (0.291) |  |  | 0.500 (0.146) |  | 35.4 | <0.001 |
| Restless Flycatcher |  |  |  |  | 0.056 (0.056) |  |  | 5.1 | ns |
| Magpie-lark | 0.400 (0.221) | 0.600 (0.267) | 0.556 (0.414) | 0.167 (0.121) | 1.167 (0.879) | 0.278 (0.135) | 0.167 (0.090) | 5.9 | ns |
| Apostlebird | 0.100 (0.100) |  | 0.611 (0.611) | 0.056 (0.056) | 0.056 (0.056) | 0.056 (0.056) | 0.111 (0.076) | 1.6 | ns |
| Jacky Winter | 0.200 (0.133) | 0.100 (0.100) |  | 0.056 (0.056) | 0.111 (0.076) | 0.278 (0.278) | 0.111 (0.076) | 4.1 | ns |
| Red-capped Robin |  |  |  |  | 0.389 (0.282) |  |  | 15.6 | <0.05 |
| Hooded Robin |  |  |  |  | 0.056 (0.056) |  |  | 5.1 | ns |
| Horsfield's Bushlark | 0.100 (0.100) | 0.400 (0.221) |  |  |  |  | 0.111 (0.111) | 18.8 | <0.01 |
| Tawny Grassbird |  |  | 0.056 (0.056) |  |  |  |  | 5.1 | ns |
| Rufous Songlark |  | 0.200 (0.133) | 1.167 (0.500) | 0.111 (0.076) | 0.056 (0.056) | 0.056 (0.056) | 0.056 (0.056) | 13.0 | <0.05 |
| Brown Songlark |  | 0.100 (0.100) |  |  |  |  |  | 10.0 | ns |
| Fairy Martin |  |  |  | 0.056 (0.056) |  |  |  | 5.1 | ns |
| Mistletoebird |  |  |  |  | 0.333 (0.114) | 0.111 (0.076) |  | 24.1 | <0.001 |
| Zebra Finch | 9.200 (5.230) | 49.800 (25.793) | 4.611 (1.932) | 0.278 (0.109) | 0.278 (0.158) | 3.944 (3.770) | 5.889 (2.593) | 34.0 | <0.001 |
| Double-barred Finch | 0.200 (0.200) | 0.400 (0.400) | 0.167 (0.121) | 0.500 (0.246) | 0.278 (0.177) | 0.667 (0.464) | 1.722 (1.446) | 2.4 | ns |
| Plum-headed Finch |  |  |  |  |  | 2.111 (2.111) |  | 5.1 | ns |
| Australasian Pipit |  | 0.600 (0.340) |  | 0.056 (0.056) |  |  |  | 23.2 | <0.001 |
| **THINNED** | | | | | | | | | |
| Emu | 0.900 (0.795) | 0.100 (0.100) |  |  |  | 1.000 (1.000) |  | 6.1 | ns |
| Common Bronzewing | 0.200 (0.133) |  | 0.100 (0.100) |  |  | 0.600 (0.600) | 0.600 (0.600) | 5.4 | ns |
| Crested Pigeon |  | 0.200 (0.133) | 0.500 (0.401) |  |  | 1.000 (0.316) | 0.200 (0.200) | 19.5 | <0.01 |
| Southern Squatter Pigeon |  |  | 0.100 (0.100) | 0.700 (0.700) |  | 0.200 (0.200) |  | 4.4 | ns |
| Diamond Dove |  | 0.400 (0.306) | 0.100 (0.100) |  |  | 0.200 (0.200) | 2.000 (1.049) | 22.3 | <0.01 |
| Peaceful Dove |  |  | 0.500 (0.342) |  | 0.200 (0.200) | 0.800 (0.200) |  | 24.1 | <0.001 |
| Tawny Frogmouth | 0.100 (0.100) |  |  |  |  |  |  | 4.5 | ns |
| Australian Owlet-nightjar |  | 0.100 (0.100) |  |  |  |  |  | 4.5 | ns |
| Black-breasted Buzzard |  |  |  |  | 0.200 (0.200) |  |  | 10.0 | ns |
| Whistling Kite |  |  |  |  |  | 0.200 (0.200) |  | 10.0 | ns |
| Black Kite |  | 0.300 (0.213) |  |  | 0.200 (0.200) |  |  | 8.3 | ns |
| Brown Goshawk |  | 0.200 (0.133) |  |  |  |  |  | 9.2 | ns |
| Collared Sparrowhawk | 0.100 (0.100) |  |  |  |  |  |  | 4.5 | ns |
| Spotted Harrier |  |  |  | 0.100 (0.100) |  |  |  | 4.5 | ns |
| Wedge-tailed Eagle | 0.100 (0.100) |  | 0.500 (0.307) | 0.200 (0.133) | 0.400 (0.245) | 0.400 (0.400) |  | 6.6 | ns |
| Nankeen Kestrel |  | 0.200 (0.200) |  |  |  |  |  | 4.5 | ns |
| Brown Falcon |  | 0.400 (0.163) | 0.100 (0.100) |  |  | 0.200 (0.200) | 0.200 (0.200) | 10.7 | ns |
| Australian Bustard |  |  |  |  | 0.200 (0.200) |  |  | 10.0 | ns |
| Red-chested Button-quail |  |  |  | 0.100 (0.100) |  |  |  | 4.5 | ns |
| Little Button-quail |  | 2.100 (0.722) |  |  |  |  | 0.200 (0.200) | 31.6 | <0.001 |
| Galah | 0.200 (0.133) | 0.300 (0.213) | 0.200 (0.133) |  | 1.000 (1.000) | 0.400 (0.245) | 0.200 (0.200) | 3.7 | ns |
| Sulphur-crested Cockatoo |  |  |  |  | 0.200 (0.200) | 0.200 (0.200) |  | 9.2 | ns |
| Cockatiel | 2.000 (0.931) | 0.200 (0.133) | 0.900 (0.458) |  |  | 1.000 (0.775) | 1.200 (0.490) | 14.4 | <0.05 |
| Rainbow Lorikeet |  |  | 0.200 (0.133) | 0.300 (0.153) | 0.800 (0.200) |  |  | 21.7 | <0.01 |
| Red-winged Parrot | 0.200 (0.200) | 1.500 (0.833) | 0.400 (0.221) | 0.400 (0.221) |  | 1.800 (0.663) |  | 13.6 | <0.05 |
| Pale-headed Rosella | 0.900 (0.482) | 0.300 (0.153) | 1.000 (0.422) | 0.200 (0.133) | 0.400 (0.245) | 0.800 (0.374) | 0.600 (0.400) | 4.1 | ns |
| Budgerigar | 0.200 (0.200) | 22.100 (14.238) |  | 0.900 (0.795) |  | 0.200 (0.200) |  | 16.8 | <0.05 |
| Channel-billed Cuckoo |  |  |  |  |  | 0.200 (0.200) |  | 10.0 | ns |
| Horsfield's Bronze-Cuckoo |  |  |  |  |  | 0.200 (0.200) |  | 10.0 | ns |
| Pallid Cuckoo |  |  | 0.300 (0.153) |  | 0.600 (0.245) |  | 0.800 (0.374) | 21.9 | <0.01 |
| Southern Boobook |  | 0.100 (0.100) |  |  |  |  |  | 4.5 | ns |
| Blue-winged Kookaburra |  |  |  | 0.100 (0.100) |  | 0.800 (0.200) | 0.800 (0.200) | 36.1 | <0.001 |
| Forest Kingfisher |  |  |  |  |  | 0.200 (0.200) |  | 10.0 | ns |
| Red-backed Kingfisher |  |  | 0.200 (0.200) |  |  | 0.400 (0.400) |  | 6.4 | ns |
| Sacred Kingfisher |  |  |  |  |  | 0.200 (0.200) |  | 10.0 | ns |
| Rainbow Bee-eater |  |  | 0.300 (0.153) |  |  | 1.200 (0.970) |  | 14.9 | <0.05 |
| Brown Treecreeper | 0.900 (0.180) | 1.100 (0.722) | 0.700 (0.213) | 1.600 (0.562) | 0.400 (0.245) | 0.600 (0.400) | 0.200 (0.200) | 8.8 | ns |
| Spotted Bowerbird |  |  | 0.600 (0.600) |  | 0.400 (0.245) | 0.800 (0.200) |  | 27.2 | <0.001 |
| Great Bowerbird |  |  |  | 0.100 (0.100) |  |  | 0.200 (0.200) | 6.4 | ns |
| Red-backed Fairy-wren |  | 0.300 (0.300) | 3.400 (2.798) | 0.700 (0.700) | 2.400 (1.470) | 1.000 (1.000) |  | 6.3 | ns |
| Variegated Fairy-wren | 2.800 (1.272) | 2.400 (1.024) | 4.300 (1.739) | 2.000 (1.022) | 4.000 (1.975) | 3.600 (1.364) | 9.400 (6.983) | 2.2 | ns |
| Weebill | 15.700 (4.536) | 7.100 (1.828) | 10.200 (2.190) | 16.700 (2.196) | 7.000 (2.168) | 7.400 (2.638) | 20.800 (3.216) | 14.3 | <0.05 |
| White-throated Gerygone |  |  |  |  |  |  | 0.200 (0.200) | 10.0 | ns |
| Yellow-rumped Thornbill | 6.300 (4.808) | 2.900 (1.169) | 1.600 (0.968) | 0.800 (0.359) | 0.600 (0.245) | 1.000 (0.548) | 0.600 (0.245) | 1.2 | ns |
| Red-browed Pardalote |  |  |  | 0.100 (0.100) |  | 0.400 (0.245) | 0.400 (0.400) | 11.6 | ns |
| Striated Pardalote | 5.000 (1.983) | 2.200 (1.153) | 3.300 (0.967) | 2.800 (0.727) | 2.400 (0.748) | 0.600 (0.400) | 1.400 (0.245) | 8.0 | ns |
| Singing Honeyeater | 1.200 (0.389) | 0.900 (0.482) | 1.800 (0.867) | 2.000 (0.494) | 2.200 (1.200) | 3.600 (1.208) | 3.800 (0.970) | 12.1 | ns |
| Grey-fronted Honeyeater | 1.200 (0.786) | 0.200 (0.133) | 0.200 (0.133) | 0.500 (0.224) | 0.200 (0.200) | 0.200 (0.200) | 0.200 (0.200) | 3.3 | ns |
| White-plumed Honeyeater |  | 0.100 (0.100) | 0.700 (0.700) |  |  |  |  | 3.6 | ns |
| Yellow-throated Miner | 3.700 (1.513) | 3.700 (1.126) | 5.200 (1.444) | 4.600 (1.500) | 1.600 (0.927) | 9.600 (4.354) | 6.600 (5.600) | 6.4 | ns |
| Spiny-cheeked Honeyeater | 0.300 (0.153) | 0.100 (0.100) | 0.500 (0.224) | 0.300 (0.153) | 0.200 (0.200) | 0.400 (0.245) | 0.600 (0.600) | 3.0 | ns |
| Rufous-throated Honeyeater |  |  |  |  |  | 1.400 (0.748) |  | 31.1 | <0.001 |
| Crimson Chat |  | 0.100 (0.100) |  |  |  |  |  | 4.5 | ns |
| Black Honeyeater |  |  |  |  | 0.200 (0.200) |  |  | 10.0 | ns |
| Brown Honeyeater |  |  |  |  | 0.200 (0.200) |  |  | 10.0 | ns |
| White-throated Honeyeater |  |  |  |  |  | 0.200 (0.200) |  | 10.0 | ns |
| Blue-faced Honeyeater |  |  |  |  |  | 0.600 (0.245) |  | 31.2 | <0.001 |
| Noisy Friarbird |  |  |  |  | 0.200 (0.200) |  |  | 10.0 | ns |
| Little Friarbird | 0.100 (0.100) |  | 2.800 (0.646) | 0.100 (0.100) |  | 1.200 (0.200) | 1.800 (0.490) | 46.2 | <0.001 |
| Striped Honeyeater | 0.300 (0.153) | 0.200 (0.133) | 0.200 (0.200) |  | 0.600 (0.400) | 1.000 (0.548) |  | 11.2 | ns |
| Grey-crowned Babbler | 0.500 (0.307) | 1.700 (1.106) | 0.400 (0.163) | 0.100 (0.100) | 1.400 (0.510) | 1.800 (1.800) | 1.400 (0.927) | 8.9 | ns |
| Varied Sittella | 0.600 (0.600) | 1.200 (0.629) | 1.200 (0.593) | 0.100 (0.100) |  |  |  | 8.6 | ns |
| Ground Cuckoo-shrike |  |  | 0.200 (0.133) |  |  |  |  | 9.2 | ns |
| Black-faced Cuckoo-shrike | 0.300 (0.153) | 0.400 (0.221) | 0.800 (0.249) | 0.200 (0.200) | 0.800 (0.374) | 0.400 (0.245) | 0.400 (0.245) | 6.7 | ns |
| White-winged Triller |  |  | 1.200 (0.554) |  |  | 0.200 (0.200) | 0.400 (0.245) | 15.3 | <0.05 |
| Rufous Whistler | 1.700 (1.055) | 1.100 (0.547) | 2.300 (0.616) | 1.100 (0.379) | 1.200 (0.490) | 1.200 (0.200) | 3.000 (1.049) | 9.9 | ns |
| Grey Shrike-thrush | 0.100 (0.100) | 0.400 (0.400) | 0.100 (0.100) | 0.400 (0.163) | 0.200 (0.200) |  | 0.400 (0.245) | 6.7 | ns |
| Crested Bellbird | 0.500 (0.167) | 0.200 (0.133) | 0.600 (0.163) | 0.600 (0.221) | 0.800 (0.200) | 0.600 (0.245) | 0.600 (0.245) | 5.8 | ns |
| Olive-backed Oriole |  |  | 0.100 (0.100) |  |  | 1.000 (0.000) | 0.400 (0.245) | 37.4 | <0.001 |
| Black-faced Woodswallow | 0.500 (0.342) | 5.700 (4.047) | 1.600 (0.718) | 0.800 (0.416) | 0.400 (0.245) |  |  | 7.5 | ns |
| Little Woodswallow |  |  | 1.600 (1.284) |  |  | 1.200 (0.970) | 0.400 (0.400) | 12.5 | ns |
| Grey Butcherbird | 0.400 (0.163) | 0.400 (0.163) | 0.500 (0.167) | 1.000 (0.149) | 0.800 (0.200) | 2.200 (0.970) | 1.000 (0.000) | 14.2 | <0.05 |
| Pied Butcherbird | 1.000 (0.471) | 1.600 (0.819) | 0.300 (0.153) | 0.300 (0.153) | 1.200 (0.200) | 0.600 (0.245) | 0.800 (0.200) | 10.0 | ns |
| Australian Magpie | 0.400 (0.221) | 0.700 (0.213) | 0.200 (0.133) | 0.500 (0.167) | 1.000 (0.000) | 0.400 (0.245) | 0.800 (0.200) | 10.9 | ns |
| Grey Fantail |  | 0.300 (0.213) | 0.400 (0.400) |  |  |  | 0.400 (0.245) | 9.5 | ns |
| Willie Wagtail | 0.600 (0.340) | 1.000 (0.422) | 1.000 (0.333) | 0.700 (0.260) |  | 2.000 (1.304) | 0.200 (0.200) | 7.5 | ns |
| Crow / Raven | 0.500 (0.167) | 0.200 (0.133) |  |  |  | 0.400 (0.245) |  | 16.0 | <0.05 |
| Torresian Crow |  |  | 0.300 (0.153) |  |  | 1.000 (0.775) |  | 14.9 | <0.05 |
| Magpie-lark | 0.400 (0.306) | 0.500 (0.307) | 0.800 (0.291) | 0.600 (0.221) | 0.200 (0.200) | 0.400 (0.245) | 0.400 (0.245) | 4.3 | ns |
| Apostlebird | 2.800 (2.691) | 1.300 (0.895) | 2.300 (2.082) | 0.100 (0.100) |  |  | 0.200 (0.200) | 3.8 | ns |
| Jacky Winter | 1.700 (0.423) | 1.400 (0.933) | 1.200 (0.512) | 1.100 (0.690) | 0.600 (0.245) | 0.400 (0.245) |  | 10.3 | ns |
| Rufous Songlark |  | 0.700 (0.496) | 0.200 (0.133) | 0.100 (0.100) |  |  | 0.200 (0.200) | 6.4 | ns |
| Mistletoebird | 0.300 (0.213) |  | 1.400 (0.476) | 0.100 (0.100) | 0.200 (0.200) | 2.600 (1.288) | 2.000 (1.049) | 22.0 | <0.01 |
| Zebra Finch | 0.500 (0.401) | 4.500 (3.541) | 1.700 (1.055) | 0.200 (0.200) |  |  |  | 8.2 | ns |
| Double-barred Finch | 0.100 (0.100) | 0.700 (0.700) | 3.100 (2.057) | 2.100 (1.110) | 0.400 (0.245) |  | 2.600 (1.166) | 10.8 | ns |
| **INTACT** | | | | | | | | | |
| Emu | 0.025 (0.025) |  |  | 0.062 (0.062) |  | 0.037 (0.037) |  | 4.0 | ns |
| Brown Quail | 0.175 (0.175) |  | 0.188 (0.187) |  |  |  |  | 4.3 | ns |
| Common Bronzewing | 0.625 (0.185) | 0.487 (0.197) | 0.719 (0.259) | 0.094 (0.052) | 0.111 (0.062) | 0.185 (0.093) | 0.296 (0.167) | 14.5 | <0.05 |
| Crested Pigeon | 0.325 (0.083) | 0.077 (0.043) | 0.156 (0.065) | 0.188 (0.105) | 0.185 (0.185) | 0.037 (0.037) |  | 20.7 | <0.01 |
| Southern Squatter Pigeon |  | 0.744 (0.669) | 0.406 (0.167) | 0.188 (0.083) |  |  |  | 22.5 | <0.01 |
| Diamond Dove |  | 0.641 (0.420) |  | 0.031 (0.031) |  |  | 0.222 (0.123) | 19.1 | <0.01 |
| Peaceful Dove | 0.500 (0.143) | 0.128 (0.066) | 0.375 (0.117) | 0.281 (0.092) | 0.704 (0.117) | 1.259 (0.204) | 0.630 (0.201) | 45.3 | <0.001 |
| Bar-shouldered Dove | 0.100 (0.048) | 0.128 (0.066) | 0.188 (0.070) | 0.219 (0.074) | 0.296 (0.090) | 0.444 (0.145) | 0.185 (0.120) | 11.0 | ns |
| Tawny Frogmouth |  | 0.077 (0.043) |  |  |  |  |  | 14.4 | <0.05 |
| Australian Owlet-nightjar | 0.300 (0.073) | 0.513 (0.103) | 0.062 (0.043) |  | 0.037 (0.037) |  | 0.037 (0.037) | 50.9 | <0.001 |
| Square-tailed Kite |  |  | 0.031 (0.031) |  |  |  |  | 6.0 | ns |
| Black-breasted Buzzard |  | 0.026 (0.026) | 0.031 (0.031) |  | 0.074 (0.074) |  |  | 4.1 | ns |
| Whistling Kite |  |  |  |  | 0.074 (0.051) |  | 0.037 (0.037) | 10.9 | ns |
| Black Kite |  | 0.103 (0.049) |  |  |  |  | 0.074 (0.051) | 15.2 | <0.05 |
| Brown Goshawk |  | 0.077 (0.043) |  |  |  | 0.037 (0.037) |  | 11.1 | ns |
| Collared Sparrowhawk |  |  |  | 0.031 (0.031) |  |  | 0.074 (0.051) | 10.5 | ns |
| Wedge-tailed Eagle | 0.050 (0.035) |  |  |  | 0.074 (0.051) | 0.037 (0.037) |  | 7.9 | ns |
| Nankeen Kestrel |  |  | 0.031 (0.031) | 0.031 (0.031) |  |  |  | 5.0 | ns |
| Brown Falcon | 0.125 (0.053) | 0.256 (0.080) | 0.062 (0.043) | 0.312 (0.122) | 0.037 (0.037) | 0.148 (0.070) | 0.556 (0.123) | 25.4 | <0.001 |
| Peregrine Falcon |  |  |  |  | 0.037 (0.037) |  |  | 7.3 | ns |
| Australian Bustard | 0.025 (0.025) | 0.026 (0.026) |  |  | 0.037 (0.037) |  |  | 3.6 | ns |
| Painted Button-quail | 0.175 (0.133) | 0.103 (0.072) |  |  |  |  |  | 7.4 | ns |
| Little Button-quail | 0.225 (0.104) | 0.333 (0.144) | 0.062 (0.062) | 0.188 (0.138) |  |  | 0.333 (0.131) | 14.8 | <0.05 |
| Red-tailed Black-Cockatoo |  |  | 0.125 (0.125) |  | 0.111 (0.111) | 0.037 (0.037) |  | 4.9 | ns |
| Galah | 0.200 (0.130) | 0.231 (0.119) | 0.125 (0.059) | 0.031 (0.031) | 0.037 (0.037) | 0.185 (0.151) | 0.074 (0.051) | 3.8 | ns |
| Sulphur-crested Cockatoo | 0.025 (0.025) | 0.026 (0.026) |  |  | 0.074 (0.051) |  | 0.074 (0.074) | 5.6 | ns |
| Cockatiel | 0.050 (0.035) | 0.846 (0.351) | 0.094 (0.069) |  |  | 0.111 (0.062) | 0.185 (0.076) | 24.5 | <0.001 |
| Rainbow Lorikeet | 0.200 (0.073) | 0.077 (0.057) | 0.062 (0.062) | 0.969 (0.244) | 2.778 (1.175) | 0.444 (0.134) | 0.444 (0.134) | 67.3 | <0.001 |
| Scaly-breasted Lorikeet | 0.075 (0.042) |  | 0.125 (0.125) |  |  |  |  | 10.4 | ns |
| Red-winged Parrot | 0.475 (0.124) | 0.487 (0.160) | 0.656 (0.204) | 0.219 (0.108) | 0.481 (0.188) | 0.259 (0.086) | 0.185 (0.093) | 6.6 | ns |
| Pale-headed Rosella | 1.025 (0.288) | 0.462 (0.164) | 0.656 (0.199) | 0.469 (0.190) | 0.630 (0.194) | 0.407 (0.134) | 0.926 (0.244) | 10.9 | ns |
| Budgerigar | 0.050 (0.035) | 6.692 (3.697) |  |  | 0.556 (0.444) | 0.074 (0.051) | 2.481 (1.648) | 30.4 | <0.001 |
| Pheasant Coucal |  |  |  | 0.031 (0.031) |  | 0.037 (0.037) |  | 5.7 | ns |
| Channel-billed Cuckoo |  |  |  |  |  | 0.037 (0.037) |  | 7.3 | ns |
| Horsfield's Bronze-Cuckoo |  |  | 0.156 (0.065) |  |  | 0.074 (0.051) |  | 23.4 | <0.001 |
| Black-eared Cuckoo |  |  |  | 0.062 (0.062) |  |  |  | 6.0 | ns |
| Shining Bronze-Cuckoo |  |  |  |  | 0.037 (0.037) | 0.037 (0.037) |  | 6.3 | ns |
| Brush Cuckoo |  |  |  | 0.031 (0.031) |  | 0.074 (0.051) |  | 10.5 | ns |
| Pallid Cuckoo | 0.050 (0.035) | 0.026 (0.026) | 0.094 (0.069) |  |  | 0.185 (0.076) |  | 16.9 | <0.01 |
| Southern Boobook |  | 0.077 (0.043) |  |  |  |  |  | 14.4 | <0.05 |
| Laughing Kookaburra |  | 0.026 (0.026) |  |  |  | 0.037 (0.037) | 0.037 (0.037) | 4.5 | ns |
| Blue-winged Kookaburra | 0.025 (0.025) | 0.026 (0.026) | 0.062 (0.043) | 0.094 (0.052) |  | 0.222 (0.082) | 0.148 (0.070) | 15.5 | <0.05 |
| Forest Kingfisher |  |  | 0.031 (0.031) |  |  |  |  | 6.0 | ns |
| Red-backed Kingfisher |  |  | 0.094 (0.069) | 0.094 (0.052) |  | 0.185 (0.093) | 0.185 (0.076) | 17.7 | <0.01 |
| Sacred Kingfisher |  |  |  |  |  | 0.185 (0.093) |  | 29.6 | <0.001 |
| Rainbow Bee-eater |  |  | 0.688 (0.385) |  | 0.037 (0.037) | 2.111 (0.418) |  | 147.7 | <0.001 |
| Dollarbird |  |  |  |  |  | 0.037 (0.037) |  | 7.3 | ns |
| Brown Treecreeper | 1.100 (0.385) | 1.949 (0.550) | 0.906 (0.306) | 1.000 (0.294) | 0.444 (0.172) | 1.074 (0.567) | 0.593 (0.263) | 5.6 | ns |
| Spotted Bowerbird | 0.400 (0.106) | 0.205 (0.111) | 0.094 (0.052) |  | 0.481 (0.154) | 0.185 (0.093) |  | 26.2 | <0.001 |
| Great Bowerbird |  |  |  | 0.062 (0.043) |  |  |  | 12.1 | ns |
| unknown Fairy-wren |  |  |  |  |  | 0.111 (0.062) | 0.074 (0.051) | 16.9 | <0.01 |
| Red-backed Fairy-wren |  |  | 0.031 (0.031) | 0.500 (0.391) | 0.963 (0.670) | 1.593 (0.597) | 0.148 (0.088) | 38.5 | <0.001 |
| Variegated Fairy-wren | 4.350 (0.965) | 2.231 (0.568) | 2.875 (0.759) | 2.188 (0.671) | 4.926 (1.117) | 4.074 (1.091) | 2.815 (1.009) | 11.2 | ns |
| Weebill | 8.525 (1.376) | 8.462 (0.885) | 6.719 (0.983) | 12.062 (1.317) | 15.333 (1.403) | 7.185 (1.177) | 10.074 (2.110) | 30.4 | <0.001 |
| Western Gerygone | 0.250 (0.069) | 0.026 (0.026) | 0.375 (0.154) | 0.094 (0.069) | 0.259 (0.101) | 0.370 (0.121) | 0.111 (0.062) | 14.7 | <0.05 |
| White-throated Gerygone | 0.050 (0.035) | 0.077 (0.057) | 0.094 (0.052) |  | 0.111 (0.062) | 0.037 (0.037) | 0.037 (0.037) | 4.8 | ns |
| Yellow-rumped Thornbill | 1.875 (0.470) | 3.077 (0.618) | 3.594 (0.747) | 1.375 (0.499) | 1.741 (0.760) | 2.852 (0.741) | 1.963 (0.422) | 15.1 | <0.05 |
| Buff-rumped Thornbill | 0.350 (0.195) |  |  |  |  |  |  | 23.4 | <0.001 |
| Inland Thornbill | 0.150 (0.067) | 0.641 (0.213) | 1.312 (0.480) | 0.281 (0.150) | 1.481 (0.563) | 0.926 (0.341) | 0.889 (0.289) | 12.5 | ns |
| Red-browed Pardalote |  | 0.077 (0.057) |  | 0.031 (0.031) | 0.111 (0.062) |  | 0.111 (0.062) | 11.3 | ns |
| Striated Pardalote | 2.950 (0.365) | 2.641 (0.548) | 1.375 (0.261) | 2.406 (0.326) | 5.148 (0.549) | 2.074 (0.471) | 2.296 (0.345) | 39.8 | <0.001 |
| Singing Honeyeater | 1.300 (0.197) | 1.667 (0.301) | 1.125 (0.245) | 1.719 (0.288) | 3.148 (0.452) | 3.444 (0.415) | 1.037 (0.146) | 48.0 | <0.001 |
| Grey-fronted Honeyeater | 5.325 (2.004) | 2.154 (0.878) | 1.688 (0.626) | 1.531 (0.564) | 1.111 (0.487) | 2.222 (0.821) | 2.593 (1.069) | 9.7 | ns |
| White-plumed Honeyeater | 0.125 (0.125) | 0.077 (0.057) |  | 0.062 (0.043) |  |  |  | 6.4 | ns |
| Yellow-throated Miner | 2.675 (0.848) | 1.615 (0.449) | 3.969 (0.761) | 3.969 (1.093) | 0.741 (0.211) | 0.667 (0.169) | 0.741 (0.211) | 34.5 | <0.001 |
| Spiny-cheeked Honeyeater | 1.600 (0.457) | 0.513 (0.151) | 0.812 (0.252) | 0.969 (0.235) | 3.630 (0.803) | 1.111 (0.222) | 0.778 (0.343) | 27.7 | <0.001 |
| Rufous-throated Honeyeater |  |  |  |  |  | 1.074 (0.486) | 0.037 (0.037) | 45.5 | <0.001 |
| Crimson Chat |  | 0.103 (0.080) |  | 0.031 (0.031) |  |  |  | 7.1 | ns |
| Brown Honeyeater | 0.050 (0.035) |  |  |  | 0.667 (0.456) |  |  | 14.8 | <0.05 |
| White-throated Honeyeater | 0.050 (0.050) |  |  |  |  |  |  | 4.6 | ns |
| Blue-faced Honeyeater | 0.125 (0.053) | 0.026 (0.026) |  |  |  |  | 0.074 (0.051) | 14.8 | <0.05 |
| Noisy Friarbird | 0.450 (0.124) | 0.051 (0.036) | 0.125 (0.059) | 0.344 (0.166) | 2.481 (1.634) | 0.296 (0.139) | 0.519 (0.247) | 16.6 | <0.05 |
| Little Friarbird | 0.975 (0.201) | 0.282 (0.137) | 2.375 (0.386) | 0.688 (0.158) | 0.259 (0.137) | 1.667 (0.374) | 0.630 (0.312) | 67.0 | <0.001 |
| Striped Honeyeater | 0.875 (0.161) | 0.564 (0.103) | 0.469 (0.168) | 0.531 (0.206) | 0.296 (0.117) | 0.407 (0.096) | 0.407 (0.110) | 13.1 | <0.05 |
| Grey-crowned Babbler | 1.500 (0.280) | 1.538 (0.414) | 1.312 (0.363) | 1.656 (0.459) | 1.000 (0.302) | 1.222 (0.568) | 1.741 (0.535) | 7.9 | ns |
| Varied Sittella | 0.825 (0.312) | 1.256 (0.397) | 1.500 (0.475) | 0.344 (0.209) | 0.815 (0.443) | 1.481 (0.401) | 0.407 (0.194) | 12.7 | <0.05 |
| Ground Cuckoo-shrike | 0.125 (0.064) | 0.026 (0.026) |  | 0.031 (0.031) |  |  |  | 11.4 | ns |
| Black-faced Cuckoo-shrike | 0.625 (0.155) | 1.385 (0.458) | 0.719 (0.197) | 0.344 (0.139) | 0.222 (0.097) | 0.370 (0.170) | 0.630 (0.170) | 11.5 | ns |
| White-bellied Cuckoo-shrike | 0.100 (0.060) | 0.128 (0.075) | 0.062 (0.062) |  | 0.074 (0.051) | 0.037 (0.037) |  | 5.3 | ns |
| White-winged Triller | 0.225 (0.127) | 0.128 (0.105) | 0.594 (0.200) | 0.031 (0.031) | 0.481 (0.326) | 0.074 (0.074) | 0.185 (0.076) | 18.5 | <0.01 |
| Rufous Whistler | 2.100 (0.352) | 2.462 (0.305) | 3.344 (0.617) | 1.750 (0.277) | 2.556 (0.326) | 4.889 (0.600) | 1.778 (0.252) | 26.8 | <0.001 |
| Grey Shrike-thrush | 0.800 (0.193) | 0.795 (0.258) | 0.594 (0.141) | 0.750 (0.201) | 0.630 (0.143) | 0.963 (0.223) | 1.333 (0.185) | 17.7 | <0.01 |
| Crested Bellbird | 1.175 (0.168) | 1.179 (0.232) | 1.062 (0.174) | 0.656 (0.115) | 1.000 (0.119) | 1.148 (0.225) | 1.111 (0.097) | 9.4 | ns |
| Olive-backed Oriole |  |  | 0.062 (0.062) |  | 0.074 (0.051) | 0.444 (0.097) | 0.074 (0.051) | 61.4 | <0.001 |
| White-breasted Woodswallow | 1.250 (0.830) | 2.103 (1.327) |  | 0.062 (0.062) |  |  |  | 24.5 | <0.001 |
| Masked Woodswallow | 0.025 (0.025) | 2.359 (1.413) |  |  |  | 0.074 (0.051) |  | 27.5 | <0.001 |
| White-browed Woodswallow | 0.075 (0.075) |  |  | 0.125 (0.059) |  |  | 0.111 (0.062) | 16.4 | <0.05 |
| Black-faced Woodswallow | 2.025 (1.014) | 1.821 (0.627) | 0.750 (0.397) | 0.688 (0.213) | 0.370 (0.132) | 0.593 (0.215) | 1.000 (0.282) | 6.6 | ns |
| Dusky Woodswallow |  | 0.026 (0.026) |  |  |  |  |  | 4.7 | ns |
| Little Woodswallow | 0.175 (0.175) | 0.308 (0.233) | 0.688 (0.252) | 0.031 (0.031) |  | 0.815 (0.333) | 0.519 (0.448) | 20.2 | <0.01 |
| Grey Butcherbird | 0.850 (0.132) | 0.744 (0.131) | 0.656 (0.106) | 1.688 (0.289) | 1.148 (0.205) | 0.778 (0.097) | 1.111 (0.145) | 19.1 | <0.01 |
| Pied Butcherbird | 0.700 (0.130) | 0.462 (0.151) | 0.594 (0.118) | 0.375 (0.087) | 0.741 (0.137) | 0.815 (0.107) | 0.963 (0.065) | 29.4 | <0.001 |
| Australian Magpie | 0.625 (0.093) | 0.410 (0.108) | 0.844 (0.246) | 0.844 (0.175) | 0.630 (0.095) | 0.259 (0.086) | 0.852 (0.103) | 19.6 | <0.01 |
| Grey Fantail | 0.900 (0.182) | 0.410 (0.126) | 1.000 (0.318) | 0.469 (0.162) | 1.407 (0.284) | 0.111 (0.062) | 1.037 (0.210) | 32.7 | <0.001 |
| Willie Wagtail | 0.400 (0.100) | 0.615 (0.145) | 0.906 (0.243) | 0.469 (0.162) | 0.296 (0.117) | 0.889 (0.317) | 0.259 (0.086) | 6.9 | ns |
| Crow / Raven | 0.150 (0.057) | 0.487 (0.137) | 0.062 (0.043) |  |  | 0.259 (0.086) |  | 34.1 | <0.001 |
| Torresian Crow | 0.150 (0.067) |  | 0.219 (0.087) |  |  | 0.148 (0.070) |  | 21.3 | <0.01 |
| Leaden Flycatcher |  | 0.051 (0.036) |  |  |  | 0.111 (0.082) |  | 10.2 | ns |
| Restless Flycatcher | 0.050 (0.050) | 0.026 (0.026) | 0.031 (0.031) | 0.031 (0.031) | 0.074 (0.074) |  |  | 1.8 | ns |
| Magpie-lark | 0.300 (0.135) | 0.256 (0.183) | 0.219 (0.098) | 0.156 (0.128) |  | 0.074 (0.051) | 0.037 (0.037) | 10.5 | ns |
| Apostlebird | 3.350 (1.759) | 2.615 (1.035) | 0.438 (0.376) | 1.688 (0.810) | 0.111 (0.111) | 0.593 (0.386) | 0.519 (0.411) | 11.8 | ns |
| Jacky Winter | 1.200 (0.287) | 1.333 (0.290) | 0.938 (0.237) | 0.719 (0.129) | 0.370 (0.121) | 0.815 (0.354) | 0.593 (0.202) | 12.5 | ns |
| Red-capped Robin |  | 0.051 (0.036) |  |  |  |  | 0.074 (0.051) | 10.2 | ns |
| Hooded Robin | 0.725 (0.376) | 0.436 (0.175) | 0.312 (0.165) | 0.031 (0.031) |  | 0.370 (0.262) | 0.074 (0.051) | 9.2 | ns |
| Rufous Songlark | 0.075 (0.055) | 0.385 (0.158) | 0.125 (0.059) | 0.312 (0.145) |  |  | 0.074 (0.051) | 15.4 | <0.05 |
| Brown Songlark |  | 0.026 (0.026) |  |  |  |  |  | 4.7 | ns |
| Silvereye |  |  |  |  | 0.074 (0.074) |  |  | 7.3 | ns |
| Mistletoebird |  |  | 0.094 (0.094) |  | 3.259 (0.467) | 0.111 (0.062) |  | 194.6 | <0.001 |
| Zebra Finch | 0.175 (0.094) | 3.692 (1.550) | 0.188 (0.105) | 0.062 (0.043) |  |  | 0.074 (0.074) | 45.5 | <0.001 |
| Double-barred Finch | 2.850 (0.833) | 6.436 (1.401) | 2.219 (0.546) | 2.406 (0.977) | 1.741 (0.612) | 2.593 (0.926) | 2.000 (0.728) | 12.5 | ns |
